# Brain Controllability and Control Energy in Gray–White Matter Fusion Network

**DOI:** 10.64898/2026.08.24.746600

**Authors:** Yuxin Liu, Kangjia Chen, Jiang Qiu, Jinpeng Niu

## Abstract

**Objective:** Brain network controllability provides a framework for understanding how structural organization shapes brain dynamics, yet current models mainly rely on white-matter connectivity and may overlook the contribution of gray-matter architecture.

**Approach:** We constructed a fusion network combining diffusion tensor imaging-derived white-matter connectivity with gray-matter morphological similarity and investigated its controllability, biological associations, heritability, phenotype prediction, and control energy.

**Main results:** Controllability derived from the fusion network preserved key topological properties of the white-matter network and was associated with neurotransmitter systems and cerebral metabolism. Compared with the white-matter connectivity-based network, fusion-based controllability showed a systematic shift toward higher heritability, improved prediction of several individual characteristics and cognitive functions, and lower modeled control energy for activating resting-state networks.

**Significance:** These findings suggest that incorporating gray-matter morphological information into a DTI-supported network provides a complementary structural representation for studying brain network controllability and state transitions. The lower control energy represents a model-derived transition cost and should not be interpreted as a direct measure of physiological energy expenditure.

## Introduction

The human brain functions as a complex network, with cognitive processes dependent on the state transitions and dynamic coordination across large-scale regions [1, 2]. Network control theory (NCT) furnishes a conceptual framework for understanding how structural connections support and constrain state transitions in the brain [3, 4]. In recent years, NCT has been increasingly employed in neuroimaging studies to identify aberrant controllability and energy inefficiencies in neurological and psychiatric disorders, including major depression [5–7], epilepsy [8, 9], schizophrenia [10], and Alzheimer’s disease [11, 12]. Importantly, it not only yields mechanistic understanding into how these disruptions arise from structural architecture but also informs the development of stimulation strategies aimed at modulating pathological dynamics [13–16].

“Controllability” refers to the role a specific brain region plays in facilitating a state transition. Recent years have witnessed growing interest in elucidating the neurobiological foundations of network controllability and its relevance to cognition and behavior. Structurally, controllability is anchored in white matter connectivity, with densely connected hub regions possessing enhanced capacity to facilitate state transitions [17]. Neurochemically, cognitive topographies can be reshaped through the engagement of different neurotransmitter systems [18]. Metabolically, reduced glucose and oxygen metabolism impairs the ability to regulate cerebral dynamics [8, 19]. Behaviorally, reduced control capacity and energy inefficiency have been associated with cognitive deficits and clinical symptoms, such as language, memory, and depressive symptom [6, 20–22]. Additionally, controllability was found to be heritable and linked to genetic risk in depression patients [23].

Nevertheless, current applications of brain-based NCT are focused on white matter structural networks derived from diffusion tensor imaging (DTI). This overlooks the possibility that local cortical characteristics may profoundly shape network-level controllability. Gray matter volume loss has been linked to the inefficient energetic control of brain dynamics [8, 24]. Therefore, damage to gray matter integrity may reduce network-level efficiency, ultimately disrupting normal state transitions. The local capacity of gray matter to process information and generate dynamic activity is shaped by their underlying microstructural architectures [25–27]. Morphological similarity networks, such as Morphometric INverse Divergence (MIND), capture individual-level coordinated microstructural organization in gray matter, reflecting cellular, molecular, and functional features embedded within cortical architecture [28]. Therefore, integrating gray matter information into the controllability model via the MIND network is expected to provide a more comprehensive description of how brain structure supports dynamic functions.

In this study, we constructed a multimodal fusion network integrating DTI-derived white matter connectivity with MIND-derived gray matter morphology and systematically investigated its controllability properties across multiple biological organizations. Specifically, we first examined whether the fusion network preserves the core topological features of controllability observed in structural networks. Next, we assessed the associations between regional controllability and neurotransmitter systems, as well as cerebral metabolic indices. We then quantified the heritability of AC to determine whether multimodal integration enhances genetic signals. Then, we evaluated the behavioral relevance of fusion-based controllability by testing its ability to predict individual differences in demographic and cognitive measures. Finally, we compared the global control energy (CE) required to activate each of the seven canonical resting-state networks under the constraints of the fusion versus DTI networks. By systematically benchmarking the fusion network against the DTI network, we aimed to determine whether multimodal integration yields a more comprehensive and functionally relevant model of brain network controllability. We found that the fusion network preserved canonical controllability patterns, increased heritability, improved behavioral prediction, and lower modeled control energy.

## Methods

### Data

#### Discovery cohort

For the discovery cohort, 354 healthy participants (28.13 ± 3.88 years, 179 females) were included from the Human Connectome Project S900 Young Adult release [29, 30]. The protocol was approved by the Washington University Institutional Review Board. All MRI data were acquired using a Siemens 3T Skyra scanner (32-channel head coil). The T1w images were acquired using a 3D MPRAGE sequence (repetition time (TR) = 2400 ms, echo time (TE) = 2.14 ms, field of view (FOV) = 224 × 224 mm^2^, matrix size = 256 × 320 × 320, flip angle = 8°, and voxel size = 0.7 × 0.7 × 0.7 mm^3^). The diffusion MRI images were acquired using a spin-echo EPI sequence (TR = 5520 ms, TE = 89.50 ms, FOV = 210 × 180 mm^2^, voxel size = 1.25 mm^3^, b value = 1000, 2,000, and 3,000 s/mm^2^, number of diffusion directions = 270, and number of b0 images = 18).

#### Replication cohort 1

For the replication cohort 1, 191 healthy participants (41.66 ± 17.16 years, 116 females) were recruited from the Southwest University Center for Brain Imaging. The protocol was approved by the Ethics Committee of Southwest University and the First Affiliated Hospital of Chongqing Medical University. All MRI images were acquired using a Siemens Trio 3.0 T scanner. The acquisition parameters for T1w images were as follows: TR = 1900 ms, TE = 2.52 ms, FOV = 256 × 256 mm^2^, matrix size = 256 × 256, flip angle = 9°, and voxel size = 1 × 1 × 1 mm^3^. The acquisition parameters for diffusion MRI images were as follows: TR = 11,000 ms, TE= 98 ms, FOV= 256 × 256 mm^2^, matrix size = 128 × 128, voxel size = 2 × 2 × 2mm^3^, slices = 60, one volume with b = 0 s/mm^2^, 30 non-collinear directions b = 1000 s/mm^2^.

#### Replication cohort 2

For the replication cohort 2, 105 healthy participants (29.11 ± 7.83 years, 61 females) were recruited from the Department of Psychiatry, First Hospital of Shanxi Medical University and the community nearby (Table S1). The protocol was approved by the local ethics committee of the Shanxi Medical University. All MRI images were acquired using a 3.0-T Trio Siemens System. The acquisition parameters for T1w images were as follows: TR = 2300 ms, TE = 2.95 ms, TI = 900 ms, flip angle = 9°, FOV = 225 × 240 mm^2^, 160 slices, thickness=1.2 mm. The acquisition parameters for diffusion MRI images were as follows: TR = 6000 ms, TE = 90 ms, flip angle = 90°, slice thickness = 3 mm, FOV = 240 mm × 240 mm^2^, matrix size = 128 × 128, voxel size = 1.875 × 1.875 × 3 mm^3^, slices = 45, one volume with b = 0 s/mm2, 12 non-collinear directions b = 1000 s/mm^2^.

This study was conducted in compliance with the Helsinki Declaration, and written informed consent was secured from all participants.

### MRI Images Pre-processing

All T1-weighted images were processed through the FreeSurfer v6.0 pipeline [31]. The sequential steps for each participant included skull stripping, intensity normalization, tissue classification (gray/white matter), volumetric labeling, surface reconstruction, and atlas registration.

Diffusion MRI images were pre-processed using the MRtrix3 pipeline (v3.0.4) [32], which included denoising, Gibbs-ringing removal, susceptibility distortion and eddy-current correction, bias field correction, and registration to T1w images. The aligned T1 image was segmented to provide anatomical constraints for tractography. To model white matter architecture, fiber orientation distributions (FODs) were estimated via multi-tissue constrained spherical deconvolution (CSD) [33]. Whole-brain probabilistic tractography was executed within the Anatomically-Constrained Tractography (ACT) framework [34], initiating 10 million streamlines using the second-order integration over FODs (iFOD2) algorithm [35]. In order to mitigate reconstruction biases, the tractogram was refined using the SIFT2 method, which optimizes streamline weights based on cross-sectional multipliers [36]. Finally, the filtered streamlines were used to construct the structural network.

### Construction of Fusion Network

To construct an individual-specific DTI network, reconstructed fiber pathways were mapped to the HCP_MMP atlas, yielding a 360 × 360 connectivity matrix in which connection strength was quantified as the number of streamlines. Consequently, the DTI network was binarized to indicate the presence or absence of inter-regional white matter tracts.

To construct individual MIND network, five vertex-wise structural features — cortical thickness, surface area, gray matter volume, mean curvature, and sulcal depth— were extracted, normalized across all vertices, and aggregated within each cortical region to form a multivariate distribution. The similarity between the multivariate distributions of each region pair was assessed using the symmetric KL divergence, estimated via a K-nearest neighbor approach. These divergence values were transformed into normalized similarity indices ranging from 0 to 1, with higher values denoting greater morphometric similarity [28]. Finally, a 360 × 360 MIND network was generated for each individual, which captures the profile similarity of multiple structural features across the cortex.

To construct an individual fusion network, the MIND network was thresholded by the binarized DTI network. Specifically, only edges present in the DTI network (indicative of direct white matter connectivity) retained their corresponding MIND similarity values (reflecting gray matter morphometric alignment), thereby creating a network that jointly captures white matter tractography connections and the biological properties of gray matter.

### Average Controllability

NCT establishes a principled framework to quantify the role of each network node in steering whole-brain state transitions. Following previous studies, we constructed a noise-free, linear time-invariant model to quantify controllability (Equation 1), where the DTI and fusion networks served as the system matrix governing brain state dynamics [37, 38].

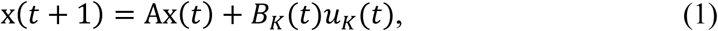

where *x* represents the brain state at specific time, matrix *A* represents the network, matrix *B_K_* describes the nodes subject to external input, and *u_K_* encodes the time-varying control strategy applied to those nodes.

Average controllability (AC) quantifies the average energy input needed to drive the brain from initial state to all possible states. Nodes with higher AC are interpreted as key drivers that facilitate state transitions with minimal control energy. Here, AC is estimated as the trace of the controllability Gramian matrix, *Trace (W_K_)*. In classic control theory, *W_K_* is defined as

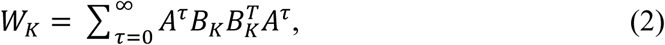

where *T* denotes the matrix transpose and *τ* represents the discrete time steps [17]. In this study, we specifically quantified and compared nodal AC between the DTI and fusion networks.

### Degree Centrality

Consistent with prior work demonstrating that regions with high AC function as network hubs and correlate with weighted degree [17], we investigated the association between regional AC and regional degree centrality in DTI and fusion networks. Degree centrality was operationalized as the average weight of edges emanating from a given region.

### Neurotransmitter Receptors and Transporters

Anatomically guided transitions across brain states can be reshaped by neurotransmitter engagement [18]. Positron emission tomography data from more than 1,200 healthy individuals were collected to quantify the distribution of 19 receptors and transporters [39]. Each neurotransmitter map was parcellated into 360 regions according to the HCP_MMP atlas. A multiple linear regression model was then employed to examine the associations between regional AC and neurotransmitter profiles. We quantified the variance explained by the model and assessed the relative contribution of each predictor using the *relaimpo* R package. Finally, we evaluated the uncertainty of the relative importance estimates using bootstrap resampling.

### Brain Metabolic Profiles

The human brain is a complex system with high metabolic demands to control brain dynamics. Following the same analytical approach used for neurotransmitter profiles, we assessed the associations between regional AC and brain metabolic profiles. Specifically, cerebral blood flow (CBF), certebral metabolic rate for oxygen (CMRO2), cerebral metabolic rate for glucose (CMRGlu), and glycolytic index (GI) were measured using positron emission tomography images in healthy individuals [40]. We then evaluated the variance explained by each metabolic profile and assessed their relative contributions to the model.

### Heritability

Heritability estimates were conducted on 327 participants in the discovery cohort, consisting of 78 MZ twins, 49 DZ twins, 175 nontwin siblings, and 25 unrelated singletons. We employed an ACE model to estimate the proportion of phenotypic variance attributable to genetic factors. By incorporating kinship information, the model partitioned the total phenotypic variance into components representing additive genetic (A), common (C) and unique environmental (E) influences. Heritability is defined as the proportion of additive genetic effects,

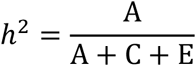

ACE model was applied to each region to generate a ROI-wise heritability estimation. To obtain network-level estimates of heritability, we aggregated the ROI-wise *h*^2^ values according to the predefined network parcellation. Moreover, to validate the influences of nuisance variables, we regressed the age, sex, age × sex, age^2^, and age^2^ × sex using a general linear model before estimating heritability.

### Support Vector Regression Analysis

We sought to determine whether the AC derived from the fusion network demonstrated superior performance in predicting individual representations, cognitive and behavioral traits. Specifically, support vector regression (SVR) models were constructed to estimate each participant’s age, body mass index (BMI), grip strength, episodic memory, intelligence, and language based on the whole-brain AC. Detailed descriptions of the motor and cognition measures are provided in the Supplementary Materials. Prediction accuracy was assessed using leave-one-out cross-validation. For each fold, feature selection and model training were conducted on the training set, followed by model evaluation on the held-out sample. Prediction performance was quantified by the correlation between predicted and observed phenotypes.

### Global Control Energy

To investigate how underlying structural architectures facilitate transitions from a baseline state to a preferential state, we employed an optimal control framework to quantify the energy required to activate specific canonical networks, including the visual, somatomotor, dorsal attention, ventral attention, limbic, frontoparietal, and default mode networks. The initial state was operationalized as a theoretical baseline with zero activity across all brain regions. For each preferential state, we set the activity magnitude of regions within a target network to one, while maintaining activity at zero in all other regions. Within this framework, an optimal solution for the control energy required at each region was derived by jointly constraining energy costs and trajectory length based on the underlying network topology. We summarized the optimal control energy globally as a measure of the brain’s energetic efficiency. For each participant, we estimated the global control energy during each transition under the constraints of structural and fusion network topology, respectively.

### Statistics and Reproducibility

All statistical analyses were performed using two-sided tests unless otherwise specified, with P < 0.05 considered statistically significant. Pearson correlation and multiple linear regression analyses were used to assess associations between controllability measures and biological features. Relative importance analysis was performed with bootstrap resampling to estimate 95% confidence intervals, and multiple comparisons were corrected using false discovery rate (FDR) correction where applicable. Heritability was estimated using an ACE model after regressing out age, sex, age × sex, age², and age² × sex effects. Support vector regression models were evaluated using cross-validation, and prediction performance was assessed by correlations between predicted and observed phenotypes. Differences between prediction models were evaluated using one-tailed Steiger’s tests; Steiger’s z values were calculated using an online implementation (https://www.psychometrica.de/correlation.html), and significance was assessed using permutation testing.

To assess the generalizability of our findings, we validated the observed patterns of AC using two independent cohorts. First, we examined whether the regional distribution of AC identified in the discovery cohort could be replicated in independent cohorts with similar imaging characteristics. Second, we replicated the association between AC in both structural and fusion networks and degree centrality across two independent cohorts. Consistent patterns of AC would provide strong evidence that the observed topological organization of control properties reflects a fundamental property of brain network architecture rather than sample-specific or protocol-specific artifacts.

## Results

### Distribution Pattern of AC

We quantified the AC of each node in both the DTI and fusion networks. AC identifies brain regions that are capable of steering the system into a diverse range of states. The DTI network and the fusion network integrating biological information exhibited similar patterns of AC (*R* = 0.92, *P*_spin_ < 0.001). Regions with higher AC included the precuneus, posterior cingulate, superior frontal, paracentral, precentral, and visual areas, whereas the medial prefrontal and temporal cortex showed relatively lower AC (Figure 1(a), (b)). The spatial distribution pattern of AC observed in the discovery cohort was reproducible across two independent replication cohorts (Supplementary Figure 1 and Supplementary Figure 2).

**Figure 1.**
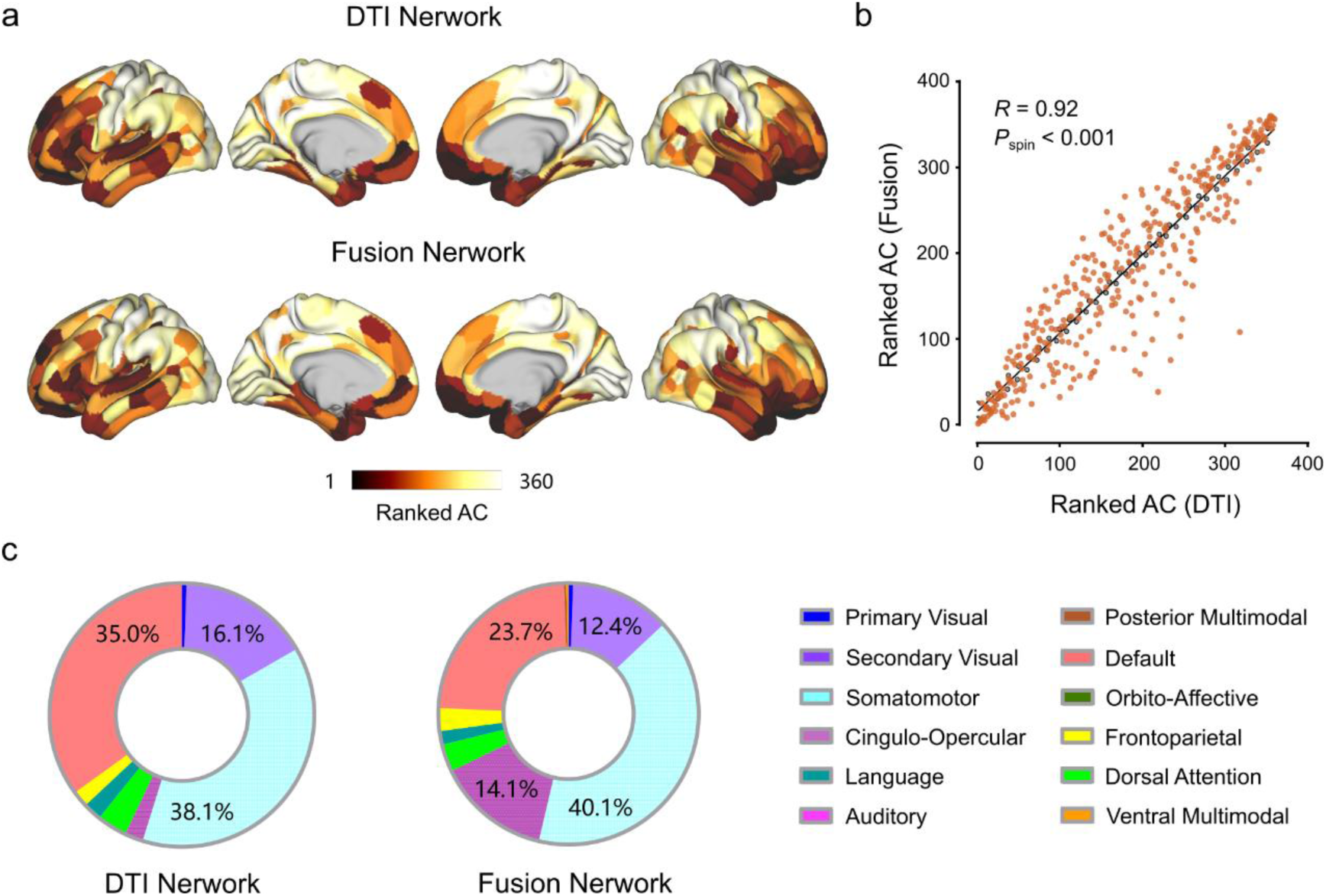
Spatial distribution of AC. **(a),** Regional AC derived from the DTI network (top row) and fusion network (bottom row) ranked on 360 brain regions. (**b),** Scatter plot of AC derived from the DTI and fusion networks. (**c),** The proportion of participants for whom the region with maximal AC fell within each network. DTI, diffusion tensor imaging; AC, average controllability.

For each participant, we identified the brain region with the highest AC and assigned it to specific network. Subsequently, we calculated the percentage of participants whose highest-AC region fell within each network, yielding a network-level distribution of controllability. We found that regions of high AC are differentially located in the somatomotor, default mode and visual systems (Figure 1(c)). Higher AC values indicate a strong capacity of the system to facilitate transitions between nearby brain states.

Importantly, AC was strongly correlated with weighted degree, suggesting that core nodes within the network also exhibit high controllability (Figure 2(a), (b)). The finding was further validated using two independent cohorts, underscoring its reliability and generalizability (Figure 2(c)–(f)).

**Figure 2.**
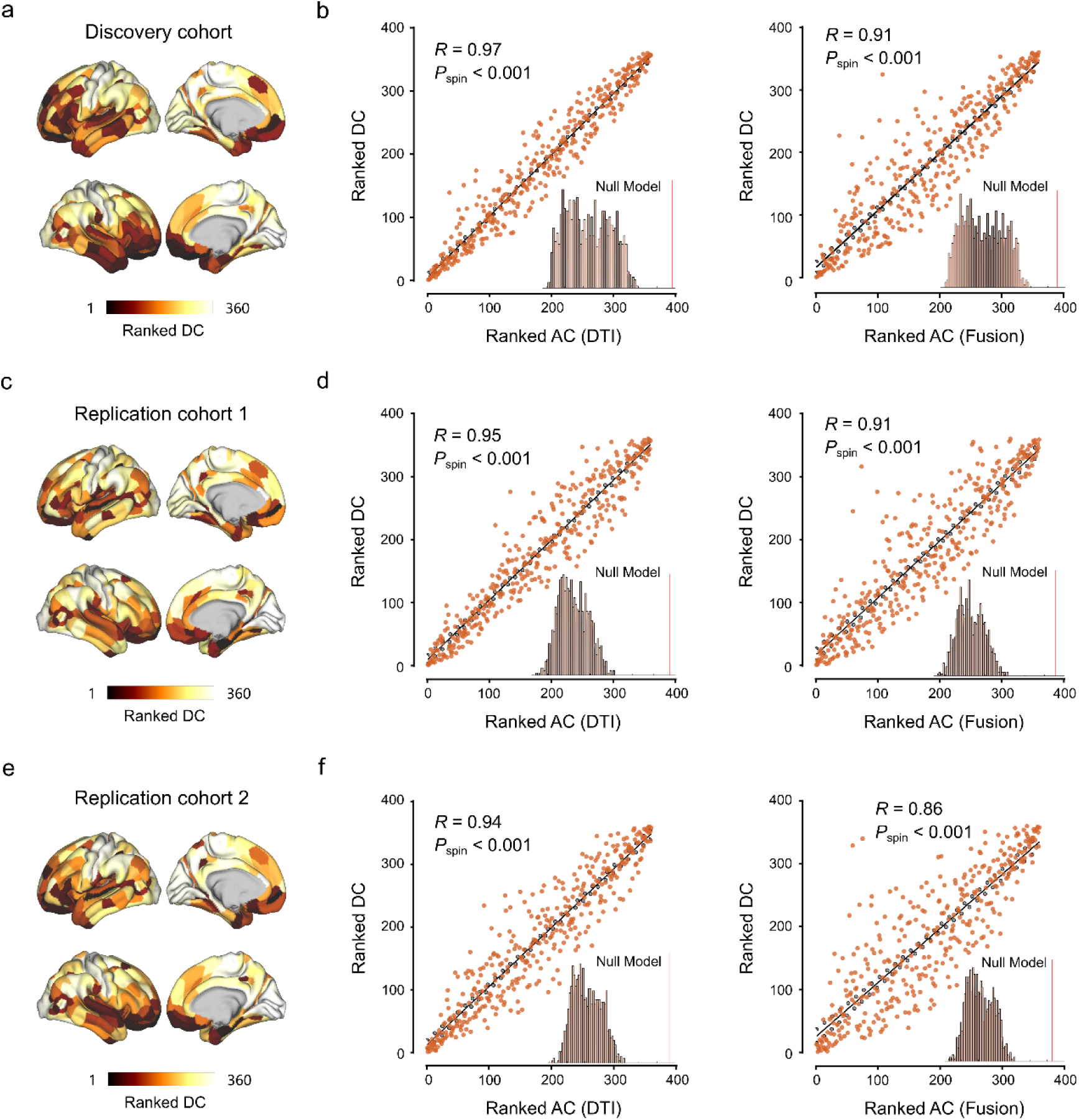
Network control properties. **(a), (c), (e),** Regional DC ranked on 360 brain regions from the discovery and replication cohorts. (**b), (d), (f),** Scatter plot of ranked weighted degree versus AC derived from the DTI network (left) and fusion network (right). AC, average controllability; DC, degree centrality.

### Association with Neurotransmitter System

To investigate the neurochemical underpinnings of network controllability, we examined the associations between regional AC and the distribution of 19 neurotransmitter receptors and transporters derived from PET imaging. For DTI network, we found that neurotransmitter profiles explained 31.83% of the variance in AC (*F*_(19, 340)_ = 8.36, *P* < 2.2×10^−16^, adjusted *R*^2^ = 0.28). For fusion network, the neurotransmitter profiles explained 32.53% of the variance in AC (*F*_(19, 340)_ = 8.63, *P* < 2.2×10^−16^, adjusted *R*^2^ = 0.29; Figure 3). These findings suggest that regions with high controllability are preferentially enriched with specific neurotransmitter systems, pointing to a potential molecular basis for the brain’s capacity to facilitate state transitions.

**Figure 3.**
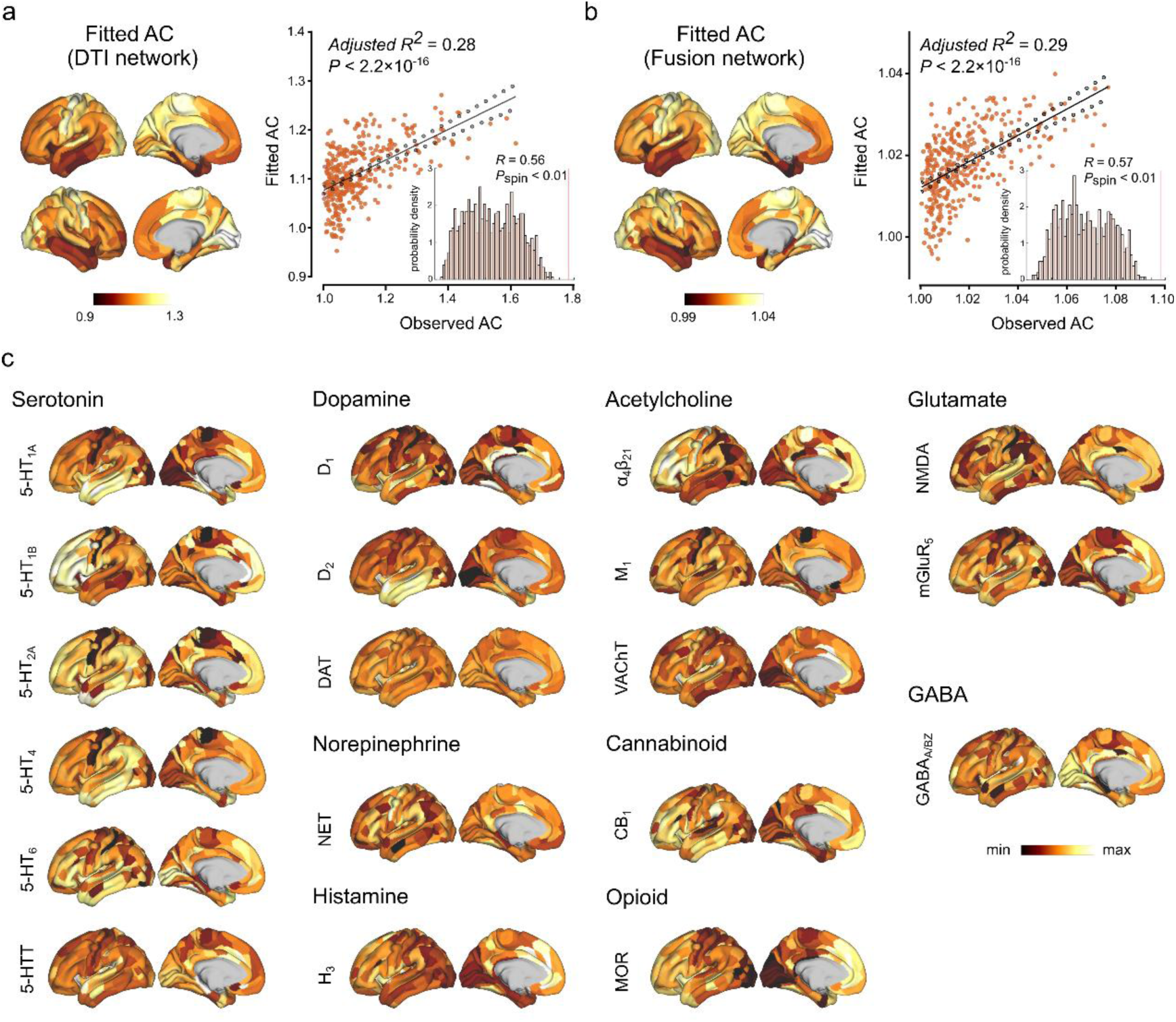
Associations with neurotransmitter. The relationship between neurotransmitter and AC derived from the DTI network (**a**) and fusion network (**b**). The brain maps illustrate the fitted AC. The scatter plots illustrate the correlation between the predicted and observed AC. (**c),** Cortical neurotransmitter density maps of 19 neurotransmitter receptors and transporters. AC, average controllability.

### Association with Metabolic Profile

We next examined the relationship between regional AC and cerebral metabolic profiles, including CBF, CMRO₂, CMRglu, and GI. Multiple linear regression analyses revealed that these metabolic indices explained 16.77% (*F*_(4, 355)_ = 17.88, *P* = 2.2×10^−13^, adjusted *R*^2^ = 0.16) and 16.74% (*F*_(4, 355)_ = 17.85, *P* = 2.3×10^−13^, adjusted *R*^2^ = 0.16) of the variance in DTI and fusion network controllability, respectively. Relative importance analysis confirmed that CMRO₂ and CMRglu were the dominant contributors (Figure 4). These findings suggest that regions with high controllability are characterized by elevated metabolic demand, consistent with their role as network hubs supporting dynamic state transitions.

**Figure 4.**
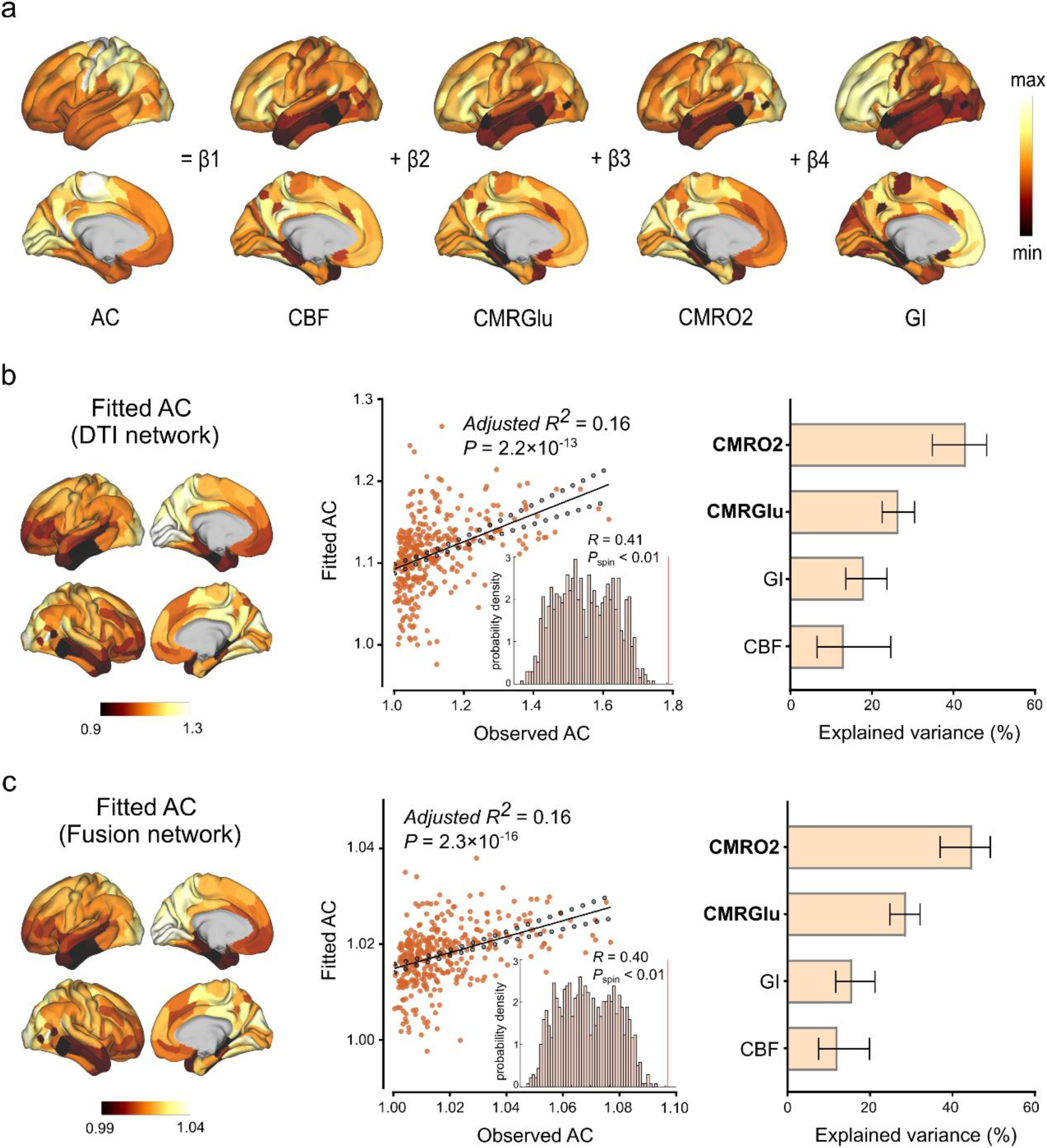
Associations with brain metabolism. **(a),** Multiple linear-regression models were used to determine the relationship between AC and brain metabolic profiles. The relationship between brain metabolism and AC derived from the DTI network (**b**) and fusion network (**c**). The brain maps illustrate the fitted AC. The scatter plots illustrate the correlation between the predicted and observed AC. The bar graphs illustrate the relative contribution of each metabolic profile for predicting the AC. Error bars represent 95% bootstrap confidence intervals, and bold font represents *P*_FDR_ < 0.05. AC, average controllability; CBF, cerebral blood flow; CMRGlu, cerebral metabolic rates for glucose; CMRO2, cerebral metabolic rates for oxygen; GI, glycolytic index.

### Heritability of AC

To investigate the genetic underpinnings of network controllability, we estimated the heritability of AC in both DTI and fusion networks. Node-level analysis revealed that heritability estimates in the DTI network were relatively high in the lateral temporal and medial prefrontal cortex (Figure 5(a)). In the fusion network, higher heritability of AC was observed across broader cortical areas (Figure 5(b)). At the network level, higher numerical heritability estimates were observed, notably in the default mode and higher-order multimodal association networks (Figure 5(c)). Finally, cumulative distribution demonstrated a systematic rightward shift in the heritability estimates for the fusion network relative to the DTI network, indicating a consistent enhancement of genetic influences across the entire distribution of regional heritability. These findings suggest that integrating multimodal biological information into the fusion network shows a systematic shift toward higher heritability estimates on brain network organization.

**Figure 5.**
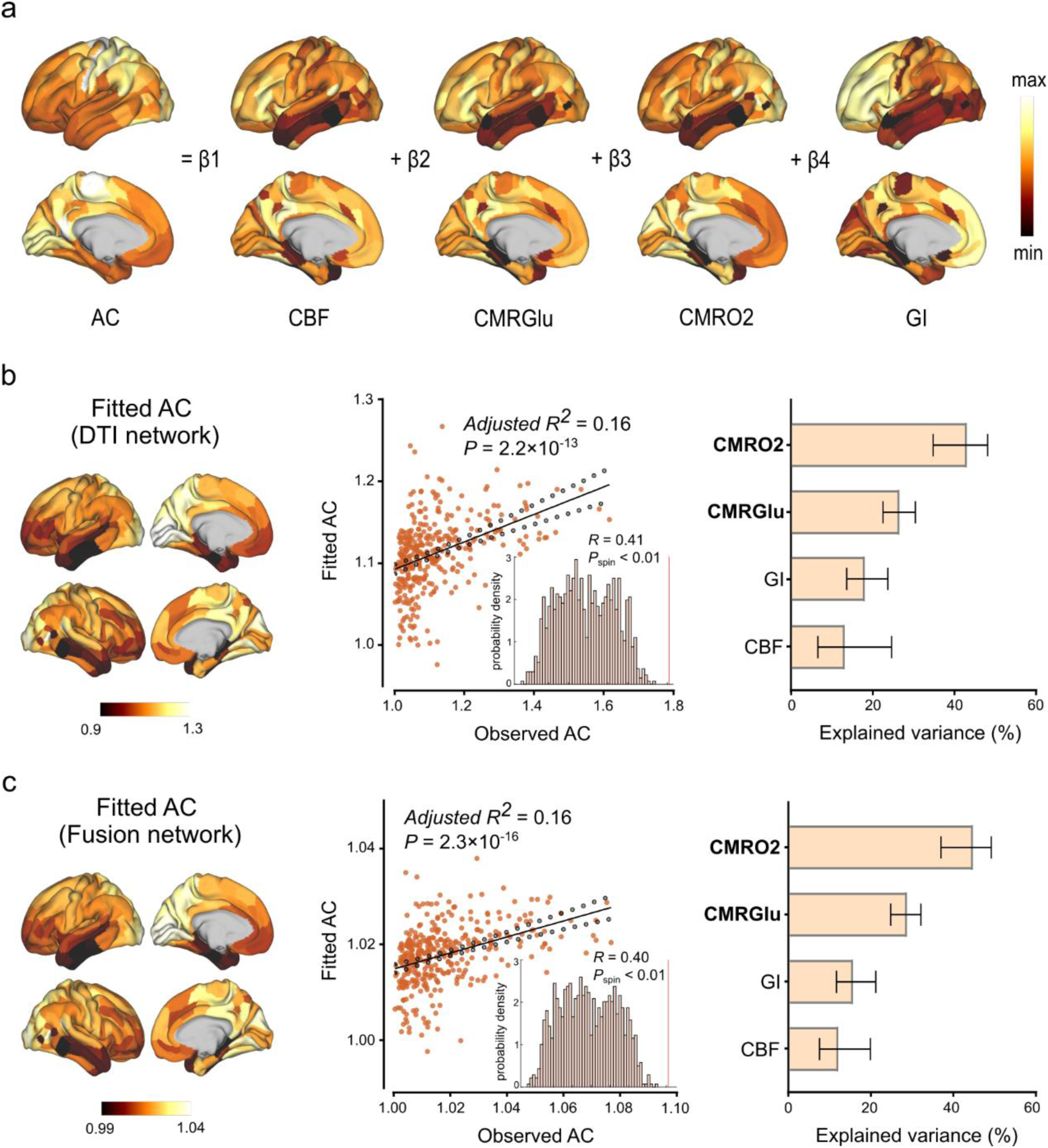
Heritability of AC. ROI-wise heritability of AC derived from the DTI network (**a**) and fusion network (**b**). **(c),** Network-wise heritability of AC.

### Prediction of Individual Phenotypes

We next compared the predictive performance of AC derived from the fusion network versus the DTI network for behavioral and demographic measures. Compared to the DTI network, AC derived from the fusion network showed superior predictive performance across multiple individual phenotypes. Specifically, the AC of the fusion network emerged as a significant predictor of both age (*R* = 0.23, *P* = 2.9×10^−5^), BMI (*R* = 0.13, *P* = 0.02), grip strength (*R* = 0.39, *P* = 1.6×10^−13^), episodic memory (*R* = 0.19, *P* = 4.1×10^−4^), intelligence (*R* = 0.22, *P* = 5.0×10^−5^), language (*R* = 0.13, *P* = 0.016), and processing speed (*R* = 0.11, *P* = 0.04) (Figure 6(a)–(g)). Direct comparisons of the dependent prediction–observation correlations using one-tailed Steiger’s tests further confirmed that AC derived from the fusion network significantly outperformed AC derived from the DTI network in predicting age (*Z* = 2.25, *P* = 0.012), BMI (*Z* = 3.41, *P* <0.001), grip strength (*Z* = 2.50, *P* = 0.006), episodic memory (*Z* = 2.96, *P* = 0.002), intelligence (*Z* = 3.67, *P* <0.001), language (*Z* = 2.95, *P* = 0.002), and showed stronger but non-significant improvement in processing speed (*Z* = 1.457, *P* = 0.073). However, no significant predictive effect of AC on emotion or working memory, in either the DTI network or the fusion network (Figure 6(h), (i)). These results demonstrate that integrating multimodal biological information into the fusion network yields controllability metrics with enhanced behavioral relevance.

**Figure 6.**
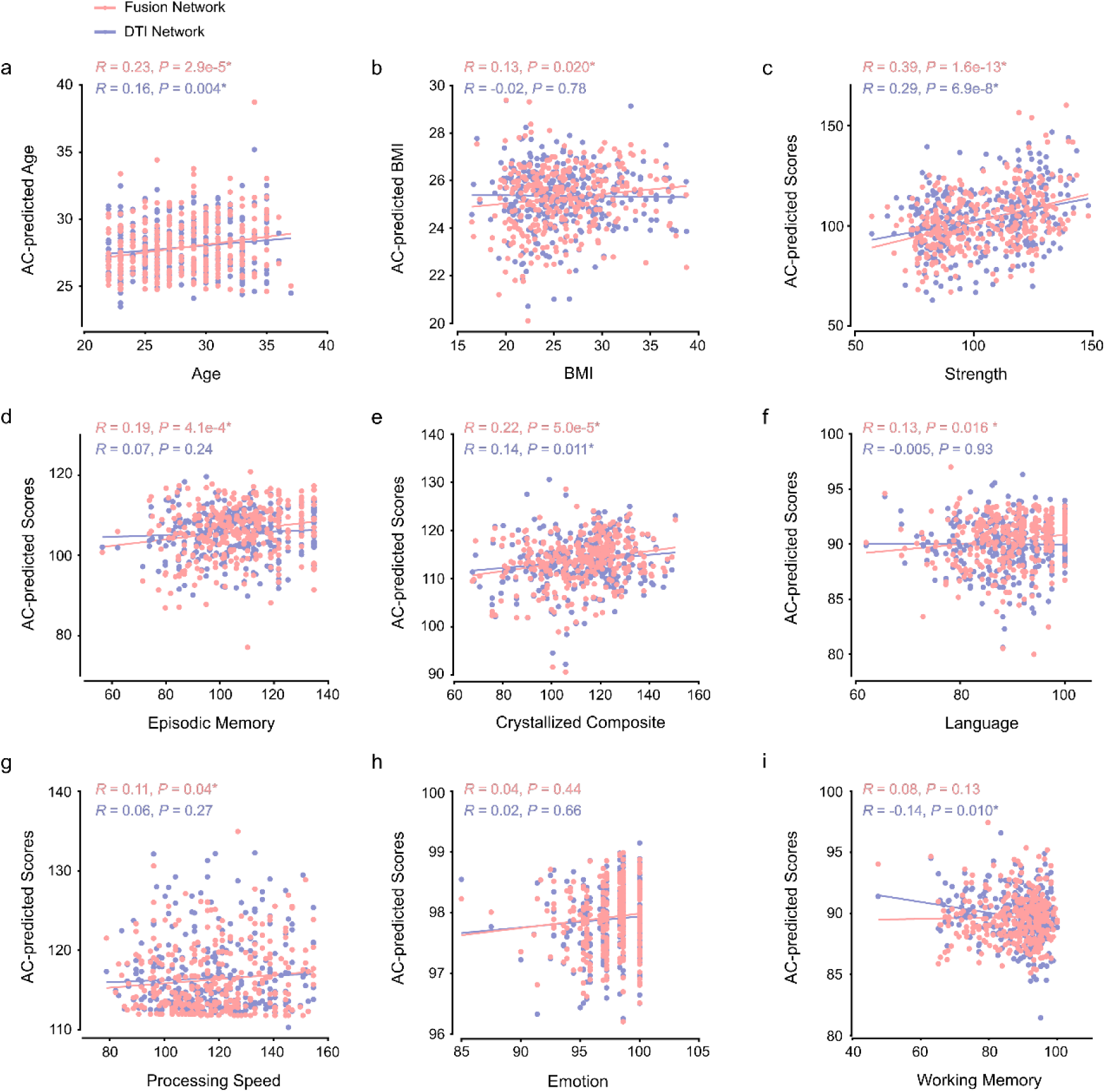
Prediction of individual phenotypes with AC. Scatter plots showing the predicted values from models based on AC compared with observed values for age (**a**), BMI (**b**), grip strength (**c**), episodic memory (**d**), intelligence (**e**), language (**f**), processing speed (**g**), emotion (**h**), and working memory (**i**). AC, average controllability; BMI, body mass index.

### Control Energy for State Activation

Using an optimal control framework, we quantified the global control energy required to activate each of the seven canonical resting-state networks from a baseline state. Comparison between network types revealed that the fusion network consistently required lower global control energy than the DTI network across all target networks (Figure 7, Supplementary Table 2). These findings suggest that multimodal data integration yields a network representation that supports more efficient modeled brain state transitions than the DTI network alone.

**Figure 7.**
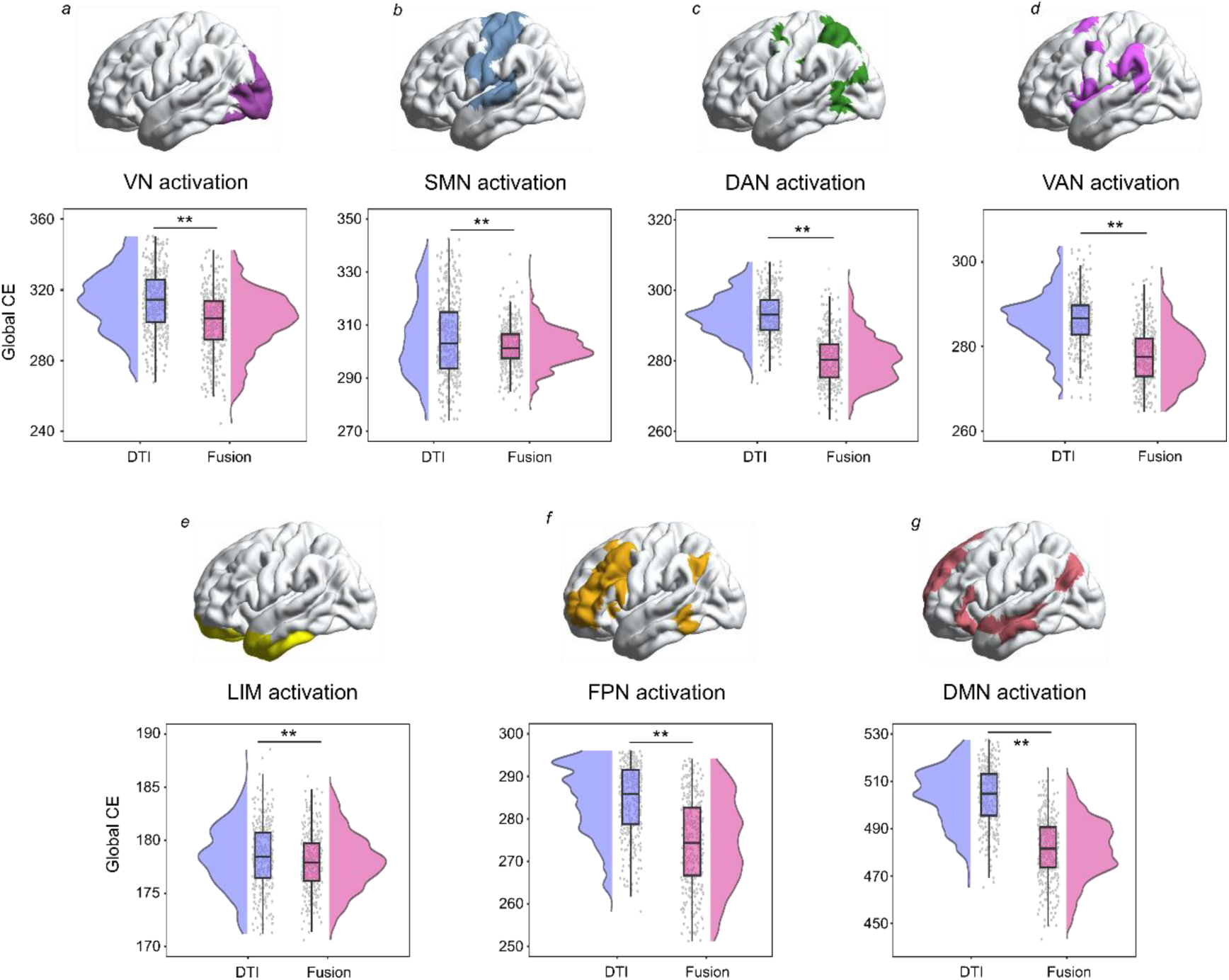
Global CE for simulating activation of canonical networks. Paired-sample t-tests were used to compare the global CE required to activate each intrinsic network under the constraints of the DTI and fusion networks. CE, control energy; VN(a), visual network; SMN(b), somatomotor network; DAN(c), dorsal attention network; VAN(d), ventral attention network; LIM(e), limbic network; FPN(f), frontoparietal network; DMN(g), default mode network.

## Discussion

Leveraging both DTI and MIND networks, we systematically evaluated the feasibility and advantages of a fusion network that integrates gray and white matter characteristics for studying brain controllability. Consistent with the DTI network, the fusion network exhibited similar spatial and hierarchical distributions of controllability, which were related to neurotransmitter systems and cerebral metabolic indices. Importantly, controllability in the fusion network exhibited both higher heritability and superior predictive power for individual phenotypes. Finally, the fusion network required lower control energy to activate intrinsic functional systems, suggesting greater energetic efficiency for dynamic state transitions. Together, these findings establish that multimodal data fusion provides a more genetically informed, neurochemically grounded, metabolically coherent, and behaviorally predictive model of brain network controllability, advancing our understanding of how structural architecture shapes brain dynamics.

Previous controllability studies have focused exclusively on DTI-derived white matter networks, they have largely ignored the potential contribution of gray matter architecture [3, 4, 24]. By incorporating MIND networks, we show that the fusion network recapitulates the fundamental controllability properties established in DTI networks. Specifically, it exhibits a comparable spatial topography, with elevated controllability in regions such as the precuneus and posterior cingulate cortex, and maintains a robust association with weighted degree. According to NCT, regions with high connection density serve as hubs that facilitate transitions to many accessible brain states, and their preferential localization within the default mode system confers higher controllability upon these regions [17, 37, 41, 42]. These findings demonstrate that while gray matter properties have been neglected in prior controllability research, their inclusion through multimodal fusion preserves the fundamental topological organization of controllability while adding biological richness, thereby validating the utility of fusion networks for studying brain dynamics.

We further confirmed the association between brain controllability and both neurotransmitter systems and cerebral metabolic profiles. Neuromodulators such as serotonin and dopamine act as general modulators of brain circuitry and play a crucial role in cognitive flexibility and state transitions [14, 39, 43, 44]. Receptors differ in their ability to promote transitions between cognitive states, and their regulatory activity is linked to cortical energy metabolism that provides the metabolic support necessary for dynamic reconfigureuration [18, 43]. Moreover, we observed the association between the controllability and oxygen and glucose metabolism. The coupling between controllability and metabolism implies that regions capable of steering the brain into many states require sustained energy supply, consistent with their role as dynamic network hubs [1, 45]. Together, these findings ground network controllability in the brain’s neurochemical environment and metabolic landscape, bridging macroscopic dynamics with molecular and physiological processes.

A key finding of this study is that controllability derived from the fusion network exhibited a systematic shift toward higher heritability than that from the DTI network alone. Gray matter morphology captured by the MIND network is highly heritable, with ample evidence demonstrating the genetic effects on cortical structure, such as thickness and volume [28, 46, 47]. Integrating these heritable gray matter features with white matter connectivity may amplify the genetic signal embedded in network controllability. Moreover, the fusion network may better approximate the biological substrate through which genetic factors shape brain dynamics, as it captures both the axonal pathways and the gray matter regions where neural computation and state transitions actually occur [27, 48]. Hence, integrating gray matter morphology with white matter connectivity captures genetically influenced aspects of brain organization that are not fully expressed in structural connectivity alone. Network controllability has recently been implicated in the genetic, individual, and familial risk architecture of major depressive disorder [49]. Therefore, the fusion network may thus serve as a more robust endophenotype for genetic association studies of controllability, offering enhanced power to detect genes influencing individual differences in brain dynamics and their perturbation in psychiatric disorders.

We found that fusion network controllability consistently outperformed DTI-based measures in predicting individual phenotypes. DTI networks primarily reflect the efficiency of structural communication pathways, which constrain but do not fully determine the brain’s dynamic repertoire [50]. Morphological similarity indexes the local computational capacity of gray matter, such as dendritic arborization, synaptic density, and neuronal integrity, which directly support cognitive processing [51, 52]. This integrative model is consistent with the observation that the larger predictive gains for cognitive phenotypes such as intelligence and language, which rely on both efficient information transfer and local processing capacity. In recent years, NCT has provided a theoretical foundation for understanding the network-level mechanisms underlying human cognitive functions and has begun to inform the development of theoretically grounded intervention strategies [45, 53, 54]. Therefore, integrating the fusion network with NCT offers a unified framework for understanding how brain structure shapes cognition and may inform future model-based intervention studies.

The human brain’s high metabolic demands and extensive connectivity necessitate control mechanisms that balance energy consumption with functional efficiency [55]. The improved energy efficiency in activating canonical networks suggests that the fusion network provides a model of brain dynamics that provides an alternative representation of modeled brain dynamics. Energy-efficient control in the brain emerges from the dynamic interaction between white matter connectivity and local cortical architecture, a synergistic relationship that becomes apparent only through their joint consideration [56, 57]. Investigations based on structural network have demonstrated energy inefficiency in neurological and psychiatric disorders, including major depressive disorder [6], epilepsy [8], and Alzheimer’s disease [11]. By integrating multimodal biological information, the fusion network may offer a more controllable architecture for detecting energy dysfunction in neurological disorders.

Several limitations should be acknowledged. First, while the fusion network integrates DTI-derived white matter connectivity with MIND-derived gray matter morphology, it does not incorporate other potentially informative modalities, such as resting-state functional connectivity or diffusion spectrum imaging metrics. Future studies could extend our framework to include additional data types for an even more comprehensive model of brain controllability. Second, the cross-sectional nature of our data precludes causal inferences about the relationships between controllability and the observed behavioral, genetic, and metabolic associations. Third, although we identified significant associations between controllability and neurotransmitter receptor systems, these findings are based on group-average receptor maps derived from independent PET samples rather than receptor densities measured in the same individuals. Finally, the age range of our sample encompasses early to middle adulthood but does not capture developmental or aging effects; whether the controllability properties of the fusion network remain stable across the lifespan or exhibit age-related changes requires investigation in broader age cohorts. Moreover, the sensitivity of the fusion network in detecting energy efficiency needs to be directly verified in the disease population.

## Conclusion

In this study, we constructed a multimodal fusion network integrating white matter connectivity with gray matter morphology to investigate brain controllability. The fusion network preserved core topological features and associations with neurotransmitter systems and cerebral metabolism. Fusion-based controllability exhibited higher heritability and better performance for predicting individual phenotypes. Moreover, the fusion network demonstrated higher efficiency in modeling transitions across brain states, supporting its value as an improved structural representation of brain controllability. Collectively, the network integrating biological information provides a more comprehensive framework for exploring how structure supports brain dynamics.

## Acknowledgements

The authors thank all participants who contributed to this study. This research received no specific grant from any funding agency in the public, commercial, or not-for-profit sectors. The authors declare that they have no financial or non-financial conflicts of interest relevant to this work.

## Data availability

The discovery dataset analyzed in this study was obtained from the Human Connectome Project (HCP) S900 Young Adult release and is available through the HCP data repository. The replication datasets were obtained from the Southwest University Center for Brain Imaging and Shanxi Medical University cohorts described in the Methods section and are available upon reasonable request to the corresponding institutions, subject to applicable ethical and data-sharing restrictions.

## Code availability

The code used for data preprocessing, construction of the gray-white matter fusion network, network controllability analysis, and statistical analyses is available from the corresponding author upon reasonable request.

## Ethics declarations

The discovery cohort was obtained from the Human Connectome Project (HCP) S900 Young Adult release. The WU-Minn HCP Consortium obtained full informed consent from all participants, and all research procedures were conducted in accordance with the Institutional Review Boards (IRB). Details of the ethical procedures are available at the HCP website (https://www.humanconnectome.org/).

The replication cohort 1 was approved by the Ethics Committee of Southwest University and the First Affiliated Hospital of Chongqing Medical University (Approval No. P12001). The replication cohort 2 was approved by the Ethics Committee of Shanxi Medical University (Approval No. HX201601). Written informed consent was obtained from all participants.

