## Supplementary Information for "Brain Controllability and Control Energy in Gray–White Matter Fusion Network"

**Motor and Cognition Measures**

Grip strength is assessed using the NIH Toolbox Grip Strength Test. Participants are instructed to squeeze a hand dynamometer as forcefully as possible, with each hand tested separately (3 minutes).

Non-verbal episodic memory was assessed using the NIH Toolbox Picture Sequence Memory Test. In this task, participants were shown a series of illustrated activities and subsequently asked to recall and reproduce the sequence in the original order. The test administration lasted approximately 7 minutes.

Language skills and crystallized cognition were assessed using two NIH Toolbox measures: Oral Reading Recognition Test and Picture Vocabulary Test. The Oral Reading Recognition Test evaluated reading decoding skills and crystallized abilities by asking participants to read aloud letters and words with maximal accuracy (4 minutes). The Picture Vocabulary Test assessed receptive vocabulary through a computer-adaptive format, in which participants selected the picture that best matched a word presented auditorily (3 minutes)

Processing speed was assessed using the NIH Toolbox Pattern Comparison Processing Speed Test. In this task, participants were shown pairs of visual stimuli and asked to determine as quickly as possible whether the two stimuli were the same or different. The test administration lasted approximately 4 minutes.

Working memory was assessed using the NIH Toolbox List Sorting Working Memory Test. In this task, participants were presented with a series of items both visually and auditorily and were asked to recall and arrange them in a specified order. The test administration lasted approximately 7 minutes

Emotion processing was assessed using the HCP Emotion Processing Task. In this task, participants were shown a target stimulus at the top of the screen and two response options at the bottom and were asked to select the option that matched the target. The task included face-matching blocks, in which the faces displayed angry or fearful expressions, and shape-matching blocks that served as a control condition. Trials were presented in blocks of six trials of the same condition, with each stimulus displayed for 2 seconds and followed by a 1-second intertrial interval. Each block was preceded by a 3-second task cue, and each of the two runs included three face blocks and three shape blocks. Overall task accuracy across the face and shape conditions was used in the present analysis.

**Supplementary Table 1. Demographic Characteristics.**

| | Discovery cohort | Replication cohort 1 | Replication cohort 2 |
|---|---|---|---|
| Number | 354 | 191 | 105 |
| Age (years) | 28.13 ± 3.88 | 41.66 ± 17.16 | 29.11 ± 7.83 |
| Sex (F/M) | 179/175 | 116/75 | 61/44 |
| BMI | 25.89 ± 4.50 | N.A. | N.A. |

Data are presented as numbers or means ± standard deviations. BMI, body mass index; N.A., not available.

**Supplementary Table 2. Statistical results of global control energy for state activation.**

| Resting-state network | t value | P value |
|---|---|---|
| Visual | -27.03 | <0.001 |
| Somatomotor | -5.76 | $1.79\times10^{-8}$ |
| Dorsal attention | -60.16 | <0.001 |
| Ventral attention | -35.36 | <0.001 |
| Limbic | -5.81 | $1.43\times10^{-8}$ |
| Frontoparietal | -40.80 | <0.001 |
| Default mode | -69.10 | <0.001 |

Paired-sample t-tests were performed to compare global control energy between fusion and DTI networks across seven canonical resting-state networks

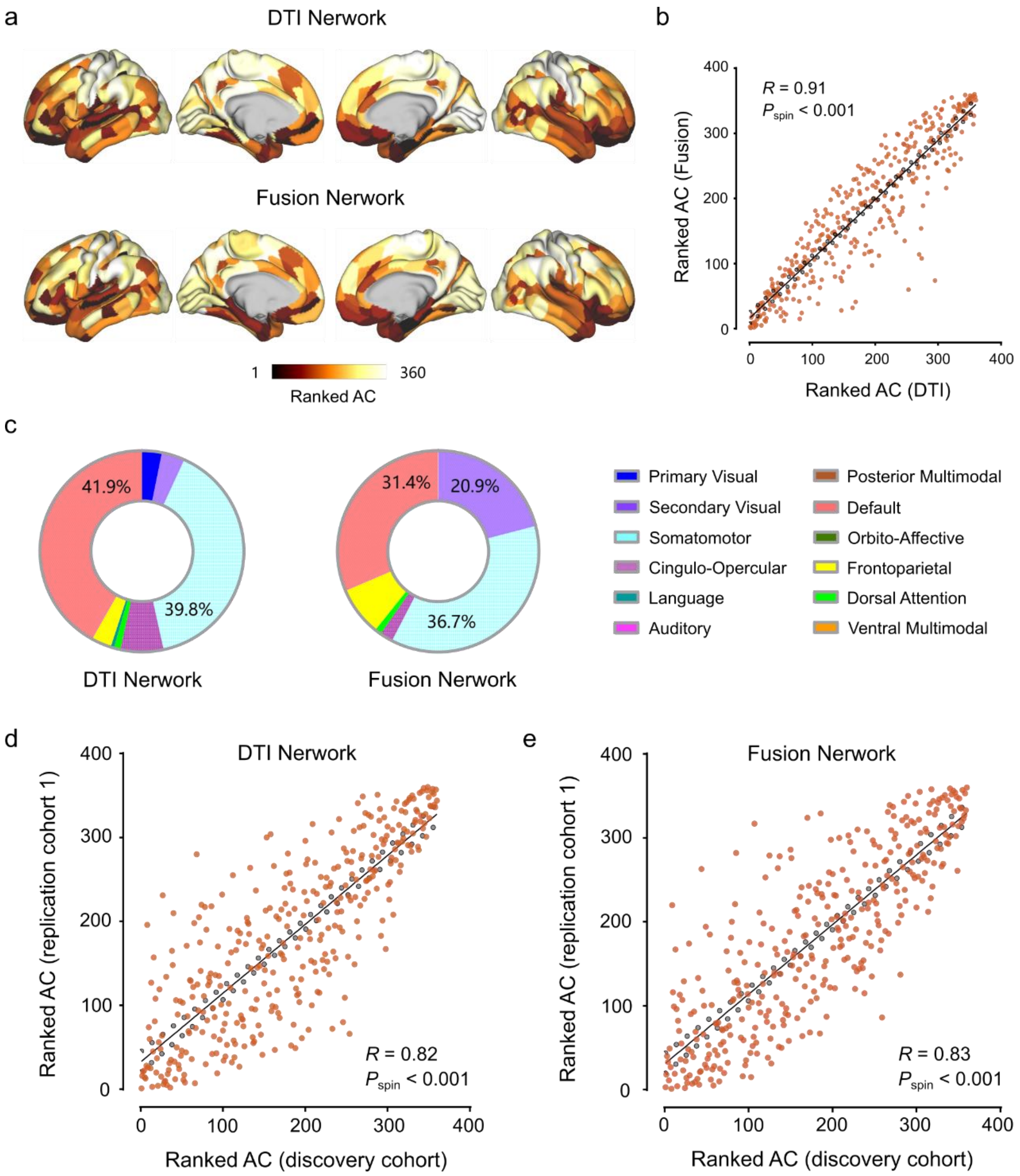


**Supplementary Figure 1 Spatial distribution of AC in the replication cohort 1. (a),** Regional AC derived from the DTI network (top row) and fusion network (bottom row) ranked on 360 brain regions. (**b**)**,** Scatter plot of AC derived from the DTI and fusion networks. (**c**)**,** The proportion of participants for whom the region with maximal AC fell within each network. Scatter plots of AC derived from the DTI network (**d**) and fusion network (**e**) between the discovery cohort and replication cohort 1. AC, average controllability.

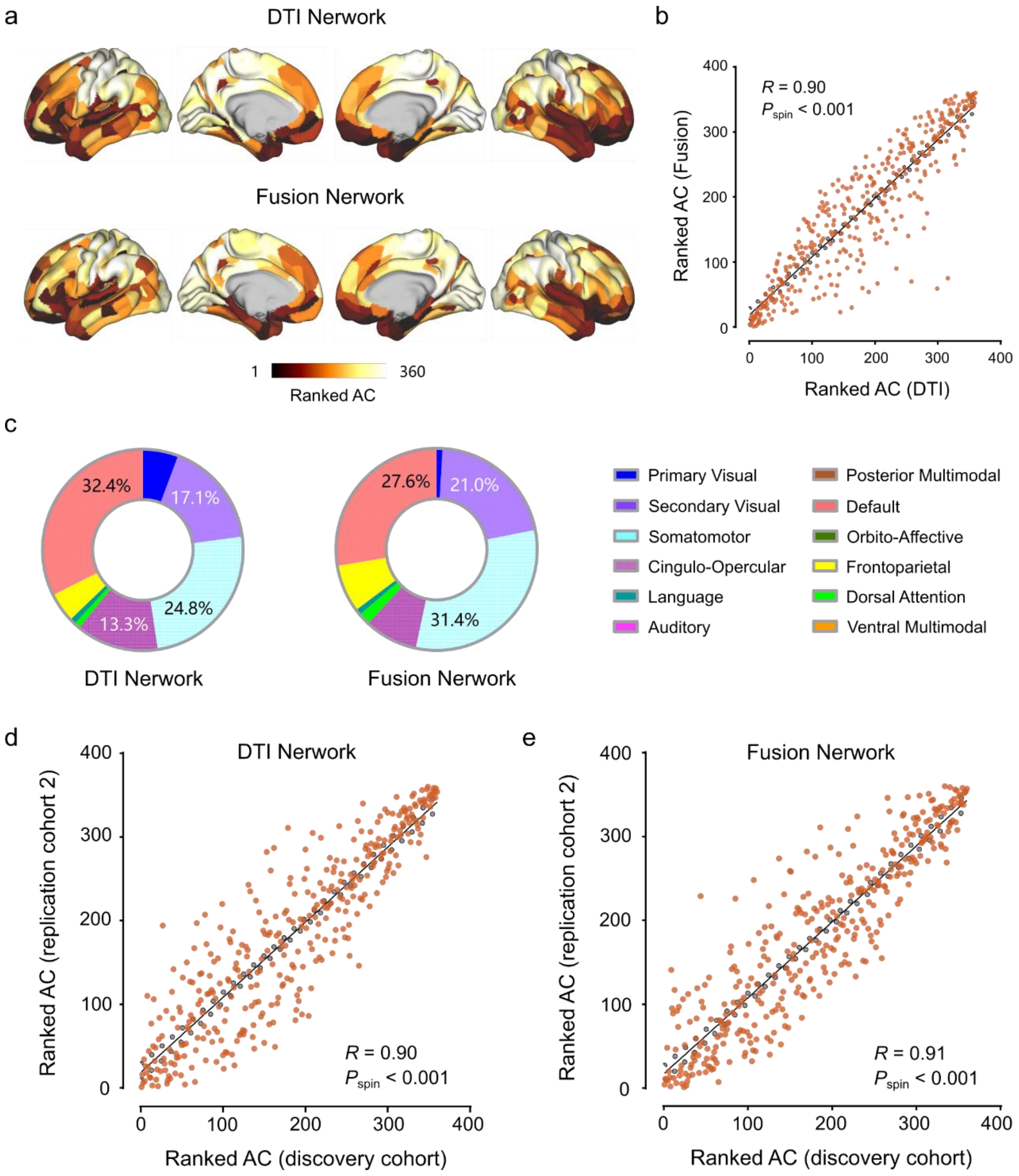


**Supplementary Figure 2 Spatial distribution of AC in the replication cohort 2. (a),** Regional AC derived from the DTI network (top row) and fusion network (bottom row) ranked on 360 brain regions. **(b),** Scatter plot of AC derived from the DTI and fusion networks. **(c),** The proportion of participants for whom the region with maximal AC fell within each network. Scatter plots of AC derived from the DTI network (**d**) and fusion network (**e**) between the discovery cohort and replication cohort 2. AC, average controllability.

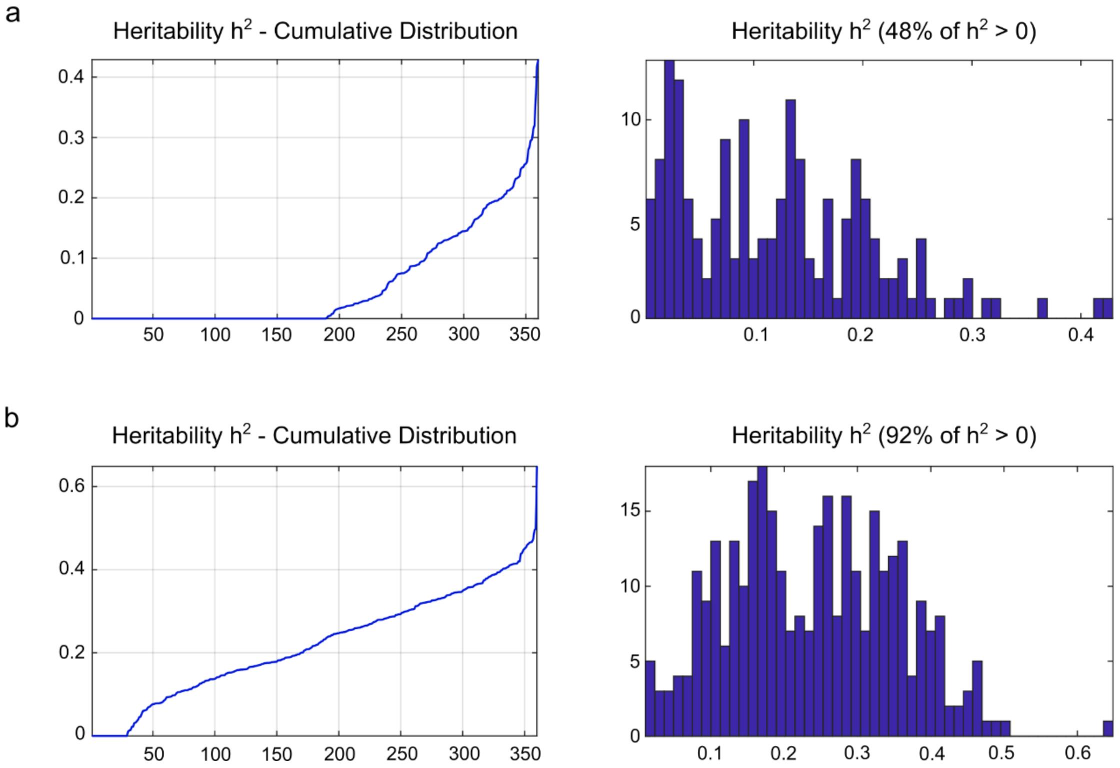


**Supplementary Figure 3 Heritability of AC. (a),** Cumulative distribution and probability distribution of the AC derived from the DTI network. **(b),** Cumulative distribution and probability distribution of the AC derived from the fusion network.
